# Calculating effect sizes in animal social network analysis

**DOI:** 10.1101/2020.05.08.084434

**Authors:** Daniel W. Franks, Michael N. Weiss, Matthew J. Silk, Robert J. Y. Perryman, Darren. P. Croft

## Abstract

1. Because of the nature of social interaction or association data, when testing hypotheses using social network data it is common for network studies to rely on permutations to control for confounding variables, and to not also control for them in the fitted statistical model. This can be a problem because it does not adjust for any bias in effect sizes generated by these confounding effects, and thus the effect sizes are not informative in the presence of counfouding variables.
2. We implemented two network simulation examples and analysed an empirical data set to demonstrate how relying solely on permutations to control for confounding variables can result in highly biased effect size estimates of animal social preferences that are uninformative when quantifying differences in behaviour.
3. Using these simulations, we show that this can sometimes even lead to effect sizes that have the wrong sign and are thus the effect size is not biologically interpretable. We demonstrate how this problem can be addressed by controlling for confounding variables in the statistical dyadic or nodal model.
4. We recommend this approach should be adopted as standard practice in the statistical analysis of animal social network data.

## Introduction

Criticism of the use of p values (Halsey *et al.* 2015; Amrhein, Greenland & McShane 2019) has arisen mainly due to researchers’ past tendency to over focus on statistical significance, rather than a balanced consideration of it along with biological importance of model effects. Consequently, there is growing movement towards the use of approaches that place greater emphasis on biological importance through effect sizes, such as model comparisons and Bayesian inference (Burnham, Anderson & Huyvaert 2011; Cumming 2014; Halsey 2019), although careful use of p values alongside effect sizes remains largely supported. Nakagawa and Cuthill highlight that “All biologists should be ultimately interested in biological importance, which may be assessed using the magnitude of an effect, but not its statistical significance” ((Nakagawa & Cuthill 2009), page 1). Correctly using effect sizes enables interpretation of the extent of an effect, allows for comparisons of the relative importance of different effects, facilitates the use of results in meta-analyses and makes it possible to use results to predict future outcomes.

While the rest of ecology has been moving towards a balanced consideration of biological significance through careful assessment of effect sizes alongside statistical significance, studies of animal social networks have tended to focus mainly on statistical significance, and neglect effect sizes. A key reason for this is that many animal social network studies focus on permutation approaches for statistical inference; permuting data and comparing model results on the observed and permuted datasets (Croft *et al.* 2011; Farine & Whitehead 2015). Network and datastream (pre-network) permutations deal with the inherent non-independence of observations related to social network data that can violate the assumptions of some conventional statistical tests. While network permutations were originally introduced to tackle non-independence (Krackhardt 1987; Krackhardt 1988), they have since been adapted in behavioural ecology to also control for a host of confounding factors inherent to observations of free-ranging individuals that influence observed associations between individuals. These include sampling effort, location, the number of times individuals are observed, and observed group sizes (Bejder, Fletcher & Brager 1998; Whitehead 2008; Croft *et al.* 2011; Farine & Whitehead 2015; Farine 2017). Indeed most processes that influence associations between individuals have typically been controlled for by imposing constraints on data permutations in these null models, for example, by constraining swaps of pairs of observations to occur only within the same time period (e.g. a day or season) or location in which they were observed.

A common procedure in network studies has been to fit a statistical model to observed network data, derive a test statistic, and then compare this test statistic to statistics derived from constrained permutations. In this procedure, confounds accounted for in the permutation are not controlled for in the fitted model. Thus, while permutations are intended to prevent these confounding effects from generating false positives or false negatives in the p-value, they do nothing to adjust for any bias in estimated effect sizes generated by these effects. As such, it is even possible for a study to suggest a significantly positive effect, yet the effect size reported for the model is negative. For example, imagine a scenario where there are multiple locations at which associations between individuals are automatically recorded over multiple sampling periods. The observed associations between individuals could be purely the result of social preferences to associate with particular other individuals. There are however, a host of additional ecological factors that could increase the probability of observing the individuals co-occurring at a given location, such as similarities in space use between individuals, uneven sex ratios, etc. (Farine *et al.* 2015b). If, in our example, individuals had a preference for associating with members of the other sex (disassortative mixing; i.e. males prefer to associate with females, and females with males) but males preferred one location and females preferred another, then it would appear in the observed network that individuals prefer to associate with others of the same sex, despite their actual social preferences being for individuals of the other sex. Permutations that control for location would show us that the observed effect sex on association rates is significantly greater than expected from permuted associations (i.e. compared to the null model) with the population been significantly disassorted. However, despite there being more disassortativity in the observed network than expected by chance, the effect size estimated by the original model would be negative and thus the opposite to the true direction of the effect. The effect size is then not biologically interpretable, and we are left to draw conclusions from the p value and direction of the effect alone, without any information on the biological importance of the effect.

Ultimately, whether this is a problem depends on the research question (Carter, Lee & Marshall 2015). If edges are intended to simply represent contacts between individuals then the importance of using approaches that control for gregariousness or location preferences may be diminished, because researchers are often more interested in quantifying emergent network structures than uncovering social preferences that drive social network structure. This would be the case for example, when studying disease or information spread (Hamede *et al.* 2009; Farine *et al.* 2015a; Silk *et al.* 2017), or the impact of social contacts on survival (Silk *et al.* 2009; Ellis *et al.* 2017) or stress (Brent *et al.* 2011). However, if the question is related to social preferences or behaviour, such as whether individuals have non-random social preferences, whether particular types of individuals prefer to associate with each other (Croft *et al.* 2009; Aplin *et al.* 2013), what type of social system is in place or how complex a social system is (Fischer *et al.* 2017; Ramos-Fernandez *et al.* 2018; Weiss *et al.* 2019), then these factors need to be controlled for. This could be relevant, for example, when addressing questions about the evolution of social preferences, such as shy individuals preferring to associate with bold individuals rather than other shy individuals. However, even where the question relates to contact networks, it might be appropriate to control for certain confounding effects (e.g. bias resulting from individuals being monitored for different periods of time or bias resulting from being more likely to observe bold individuals over shy).

Many questions in animal behaviour focus on the level of either dyads or nodes. The basic idea behind dyadic models is that the outcome, such as the probability of associating, is affected by the characteristics of both individuals in the dyad. Dyadic regressions – where the analysis is at the level of connections between pairs of individuals - are used less in animal social network analysis than in other fields. In particular other fields have developed methods for dyadic analysis, termed Actor–Partner Interdependence Models, which tackle the problem of interdependence without the use of or need for permutations (Kenny 1996; Chow, Claxton & van Dulmen 2015; Garcia, Kenny & Ledermann 2015; Kenny 2018). In these models, controls can be included in the statistical model that examines predictors of the strength of connection between the two individuals or predictors of node-level network measures. An example of this is shown in a paper using dyadic regression that controls for new confounding factors and debunks a previous network analysis suggesting that obesity is socially transmitted (Cohen-Cole & Fletcher 2008). Other methods such as Exponential Random Graph Models (ERGMs) (Silk & Fisher 2017), latent space models (Hoff, Raftery & Handcock 2002), and Stochastic Actor-based Models (Ilany, Booms & Holekamp 2015; Fisher *et al.* 2017) can also be used for dyadic analysis.

For questions relating to any network-level statistics that rely on indirect connections (Brent 2015), however, such as path length or clustering, then we may need to control for these factors in the association index itself. This is commonly done for overall gregariousness (Godde *et al.* 2013), but not for other key confounding factors. An underused method is to produce generalized affiliation indices (GAIs), which are produced from the residuals of a regression of observed contacts on confounding variables (Whitehead & James 2015). GAIs are promising, although the biological meaning of these association indices can be difficult to interpret, and with values typically being both positive and negative it may be difficult to apply many traditional network analysis approaches that ecologists are familiar with. In particular, most weighted network statistics such as clustering coefficient and path length assume that edge weights are strictly positive (Newman 2004).

Here we focus on dyadic and nodal analysis and use two network simulation examples to demonstrate that relying solely on permutations to control for confounding variables can result in incorrect and biased effect size estimates of animal social preferences that are not biologically meaningful. Our second simulation, in particular, represents a common scenario where individuals are sampled in different locations. We show with our examples that this problem can best be addressed by controlling for confounding variables in the statistical model, before demonstrating that this approach remains effective when analysing an example using real-world data.

## Methods

The code written in R 3.6.3 (R Core Team 2018) for both simulated examples is available in the online supporting information. We have chosen two example simulations with each illustrating the effect of a different confounding factor. The first simulation captures observation bias and the second captures social preference along with location preference. Our simulation framework is inspired by those of Whitehead and James (2015), Farine & Whitehead (2015) and Farine (2017). We also demonstrate the method using a real-world example, with social data on reef manta rays (*Mobula alfredi*) from Perryman *et al.* (2019).

### Simulation 1: Sex differences in gregariousness and conspicuousness

We designed our first model to simulate a scenario in which a study is testing for a sex difference in gregariousness - a nodal analysis. We simulated a population of 100 individuals, with an equal sex ratio, in which females were more gregarious than males, but males were easier to observe. All individuals were assigned a level of gregariousness *g* from a truncated normal distribution (bounds 0 and 1) with a mean of *G*_*F*_ for females and *G*_*M*_ for males, and standard deviation of 0.1. Individuals were also assigned a sighting frequency s from the same distribution but with mean *S*_*F*_ for females and *S*_*M*_ for males. Females are assumed to be more gregarious and thus were assigned a higher gregariousness than males. Females were also assumed to be more difficult to observe than males (perhaps due to conspicuousness, size, or behaviour) and thus are assigned a lower sighting probability than males. We systematically varied the values of *G*_*F*_, *S*_*F*_ and *S*_*M*_.

Using the *rgraph* function in the *R* package *sna* (Butts 2016), we then simulated t=100 sampling periods. In each sampling period, every individual was always observed so that we had a reference case of perfect sampling for each run. The probability that any two observed individuals were observed associating was equal to the product of their gregariousness scores. Once sampling was complete, we generated the association network using the Simple Ratio Index (SRI) (Cairns & Schwager 1987) and measured the sum of the strength of connections (weighted degree) for each individual as a measure of their observed gregariousness. For each run we then simulated sampling on the network, where each individual was observed with probability *S*_*M*_ for males and *S*_*F*_ for females. We do this by removing all associations involving individuals that were not sighted in a given sampling period.

We tested for a difference in strength between females and males using two different approaches. We expected females to have higher strength than males as a result of their higher gregariousness. First, we fitted a linear model to test the hypothesis that females have a higher strength than males: 

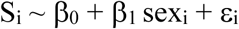

Second, we also fitted the same model but included the mean-centred number of observations as a covariate to control for sampling bias. As such we also included an interaction of observations with sex as a covariate to control for sampling bias in the effect size: 

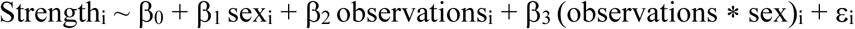

In both models we allowed unequal variance by sex. Sex was also included as an interaction with the mean-centred number of observations. This improved the model fit due to an interaction between gregariousness and sightings: because we know the sexes differ in their gregariousness we would also expect that they would have different relationships between sightings frequency and strength. For a fair comparison across parameterisations of the model, the model then compares the effect of sex at the average level of sightings frequency, and we take the ratio there as our effect size. We found that a linear relationship with the number of observations was a good fit for this model.

We used a permutation procedure to determine the statistical significance of regression coefficients for the corrected nodal model. Although our study is about effect sizes, our intention is to demonstrate their use along with permutations. Recent work has highlighted problems with using datastream permutations with standard regression to test hypotheses such as the one we are interested here (Weiss *et al.* 2020). Thus, we instead use the double semi-partialling method outlined by (Dekker et al., 2007) with 10,000 permutations, using the pivotal statistics (*t* for quasipoisson, *z* for beta) as our test statistics. This permutation procedure does not account for sampling or other confounding effects; but it does not need to, because we have already controlled for them in our statistical model.

### Simulation 2: Sex assortment and location preferences

We designed the second simulation to test the hypothesis that individuals of the opposite sex have stronger social preferences for each other than they do for individuals of the same sex. However, we also included a sex difference in location preference, that confounded the true social preferences of individuals - a dyadic analysis. We simulated 100 individuals, half of which were female and half of which were male. For all dyads of the same sex we assigned an association preference from a truncated normal distribution with a mean *S*_*Same*_=0.25 and standard deviation of 0.1 (boundaries 0 and 1). We did the same for all pairs of individuals of a different sex, but used a mean of *S*_*Different*_=0.5. This association matrix was assumed to be symmetric. We assumed two sampling locations A and B and assigned a preference for location B to each individual, using a truncated normal distribution with a standard deviation of 0.1 and means of *L*_*M*_=0.8 for males (strong preference for Location B) and *L*_*F*_=0.2 for females. To examine the effect size without bias in location preferences, we also ran the simulation with a mean preference of *L*_*M*_= *L*_*F*_=0.5 for location B (i.e. no preference) for both sexes for each set of parameter values for *S*_*same*_ and *S*_*different*_.

We then simulated t=100 sampling periods as before. In each sampling period, we assigned each individual to a location (A or B) according to their location preference. Individuals could then only associate if they were in the same location in that time point, and the probability that any two observed individuals were observed associating was proportional to the product of their association preferences. In each sampling census we recorded whether each pair of individuals were observed associating.

We again used three different ways to test for negative assortativity by sex, calculating statistical significance by comparison to the results from the permutations. First, we fitted a binomial multiple membership GLMM with logit link as a dyadic regression to test the hypothesis that individuals of the opposite sex have a higher social preference for each other than they do for individuals of the same sex. We used default priors and the model: 

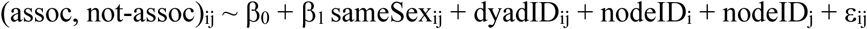

Where nodeID is a random factor in the multiple membership model. This represents a random gregariousness or sociality effect or, in the case of uneven detection, a visibility effect for each individual. Note that including nodeIDs as standard random factors without the multiple membership model would represent a directed network scenario, but we are dealing with undirected associations here. MCMCglmm includes an observation-level random intercept as default and DyadID is therefore implicitly included as a random intercept. This glmm structure follows some aspects of implementations of th Actor-Partner Interdependence Model (Kenny 1996; Kenny 2018) and the multiple membership approach is inspired by (Rushmore *et al.* 2013).

Second, we fitted the same model, but included the proportion of censuses that the two individuals in the dyad were observed in the same location (which we centred on 0.5), as a covariate to control this non-social influence: 

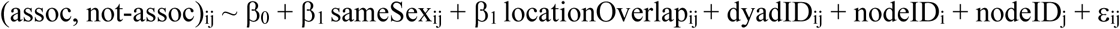

We used the multiple regression quadratic assignment procedure (MRQAP) to calculate significance for this corrected model. There are caveats with this approach. First, the GLMQAP deals specifically with the non-independence between edges that arises because they are connected to the same node, but this is already dealt with completely by the multi-membership random effect. As such, there is no need to use permutations with this approach. Additionally, MCMCglmm is a Bayesian approach, and thus p values are not a standard aspect of the analysis. However, we have included the permutation procedure here to demonstrate an appropriate permutation approach, should it be desired. MRQAP uses the double semi-partialling permutation method (Dekker et al., 2007) with 10,000 permutations. This method is equivalent to the multiple regression quadratic assignment procedure (MRQAP), but fitting GLMs instead of least squares regression. This GLMQAP procedure is available in the “aninet” R package, accessible through GitHub (https://github.com/MNWeiss/aninet).

### Reef Manta Ray Data: Sex differences in gregariousness

Simulation is the best approach for testing our method, because it allows us to know what the effect size should actually be. However, we also demonstrate the method on complex data. As such, we used group-based association data on reef manta rays (for full details see (Perryman *et al.* 2019)). Individual reef manta rays were identified by standard photo-ID methods, and data on group compositions were collected from November 2013 to May 2018 in the Dampier Strait region of Raja Ampat, West Papua, by trained researchers diving using SCUBA equipment, or freediving, depending on the position of rays in the water column. In line with (Perryman *et al.* 2019) we removed individuals observed fewer than 10 times to improve data reliability (Whitehead 2008) and derived the SRI using a 15 day sampling period. In the resulting dataset there are 70 unique females, 42 unique males, 1257 female sightings, 643 male sightings, 17.96 mean sightings per female, and 15.31 mean sightings per male. As such, there is a clear case of more females or oversampling of females.

We fitted a regression model to the network to test the hypothesis that one sex has higher strength than the other: 

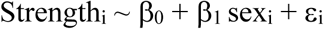

However, we know that this data is confounded by a bias towards observing females, which can bias the effect size in this respect. We also fitted the same model but included the mean-centred number of observations of each individual as a covariate to control for sampling bias in the effect size following the approach of our first simulation: 

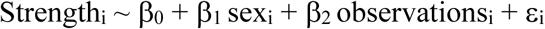

We checked the fit of observations, and a linear relationship was appropriate in this case. Unlike our nodal simulation, we did not interact sex with observations because there was no difference between the sexes in how the observations impacted strength, and it did not improve the model fit.

## Results

### Simulation 1: Sex differences in gregariousness and conspicuousness

The simulation created the desired effect that females are more gregarious than males, while males appear to be more gregarious when females are difficult to observe (Figure 1). Over 20 independent simulation runs we found that the mean effect size of sex on strength was 29.16 (95% CI = 28.66, 29.65) when sampling was perfect and males and females were equally observable (equivalent to *S*_*F*_*=*1, *S*_*M*_*=*1).

**Figure 1:**
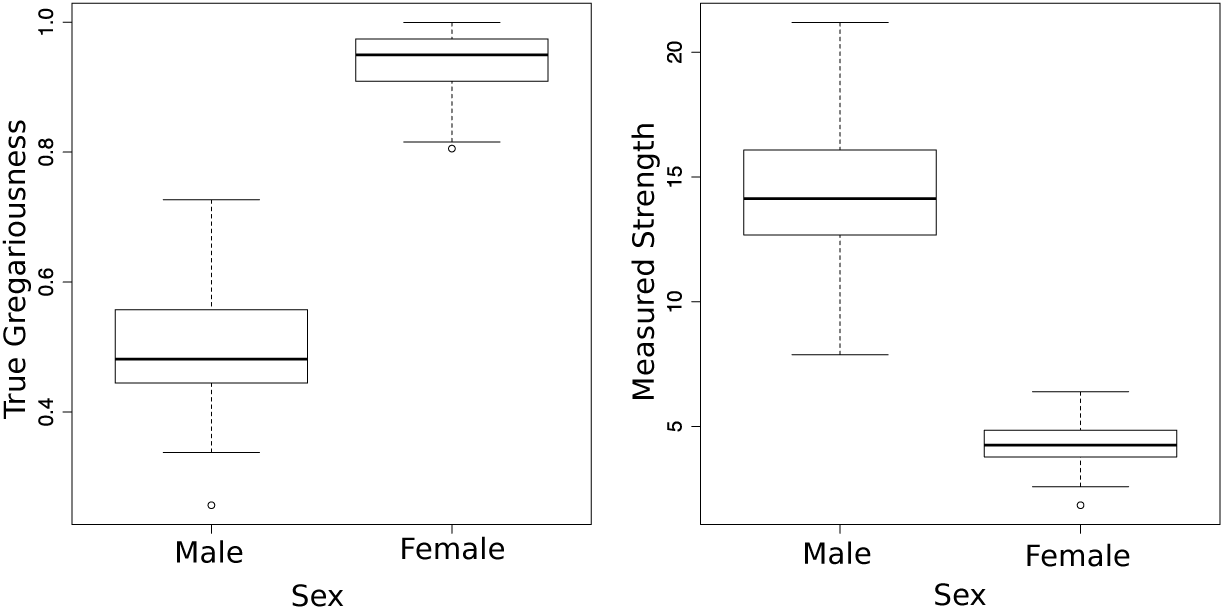
(A) The true gregariousness of individuals of each sex. (B) The measured strength of individuals of each sex from simulated observations where *S*_*M*_=1 and *S*_*F*_=0.1. In both cases *G*_*F*_=1 and *G*_*M*_=0.5. Plots are for a single representative run of the simulation. The contrast between the two panels (A) and (B) demonstrates that although females are more gregariousness, males incorrectly appear to be more gregarious when females are difficult to observe. For all box-and-whisker plots, the bottom and top of the box are the 25th and 75th percentile, the middle band is the median, the upper whisker is Q3 + 1.5IQR, the bottom whisker is Q1 + 1.5IQR, and open circles show outliers.

For each simulation run our expected effect size is from the perfectly sampled network under the given parameter values and the other two effect sizes are from the sampled network with the statistical model not accounting for observations and with the statistical model accounting for observations. Figure 2 shows the results of our analysis over systematically varied parameters *S*_*F*_ and *S*_*M*_, *where S*_*M*_>*S*_*F*_ and *S*_*M*_+*S*_*F*_=1 to keep sampling observations constant (figure 2a), and also varying *G*_*F*_ (*G*_*F*_ > *G*_*M*_) (figure 2b). The results show that not accounting for confounding effects in the statistical model (and instead relying on permutations to account for them) produces effect sizes that are very different from the actual effect sizes and for a wide range of conditions. Indeed this approach actually shows the opposite result to the reference model, in that females are shown to be less gregarious than males (points below the dotted line om figure 2a). However, when we account for sampling in the statistical model, the results are extremely similar to the actual effect sizes.

**Figure 2:**
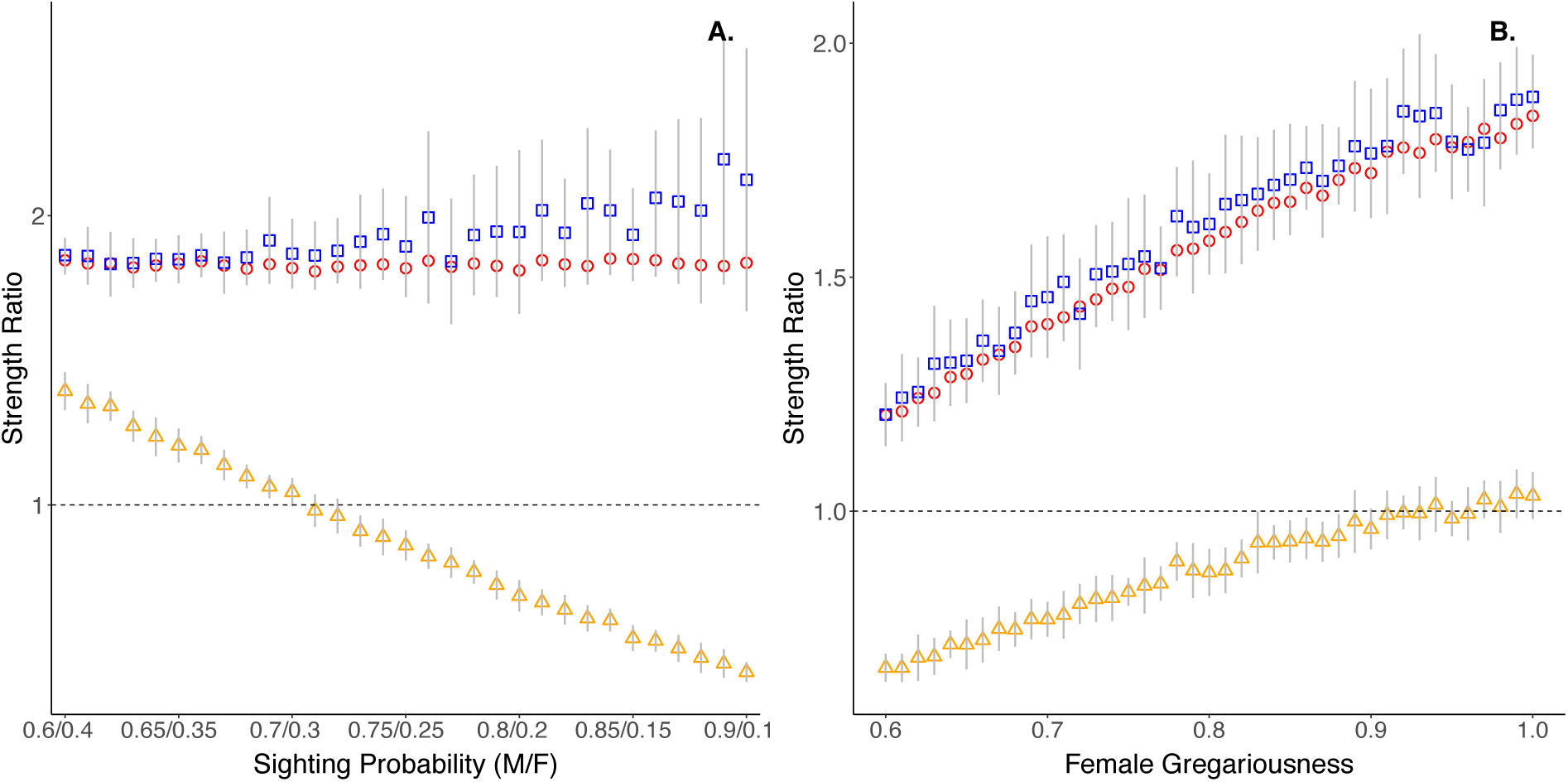
The sex effect size (ratio of strength between males and females) from the statistical models. Odds over 1 (dotted line) mean that females are more gregarious and have a higher strength than males. Red circles show the true effect size, orange triangles show the effect size of the uncorrected statistical model after biased sampling, and blue squares show the effect size of the model statistically controlling for observation bias after biased sampling. Error bars show standard deviation. Panel A shows results when G_F_ = 1.0 and G_M_ = 0.5 while covarying *S*_*F*_ *and S*_*M*_, and panel B shows results when *S*_*F*_*=0.3, S*_*M*_ = 0.7, and G_M_=0.5 while varying G_F_. These figures show that failing to account for the bias in the statistical model gives incorrect effect sizes, and that this problem can be addressed by controlling for bias in the statistical model.

**Figure 3:**
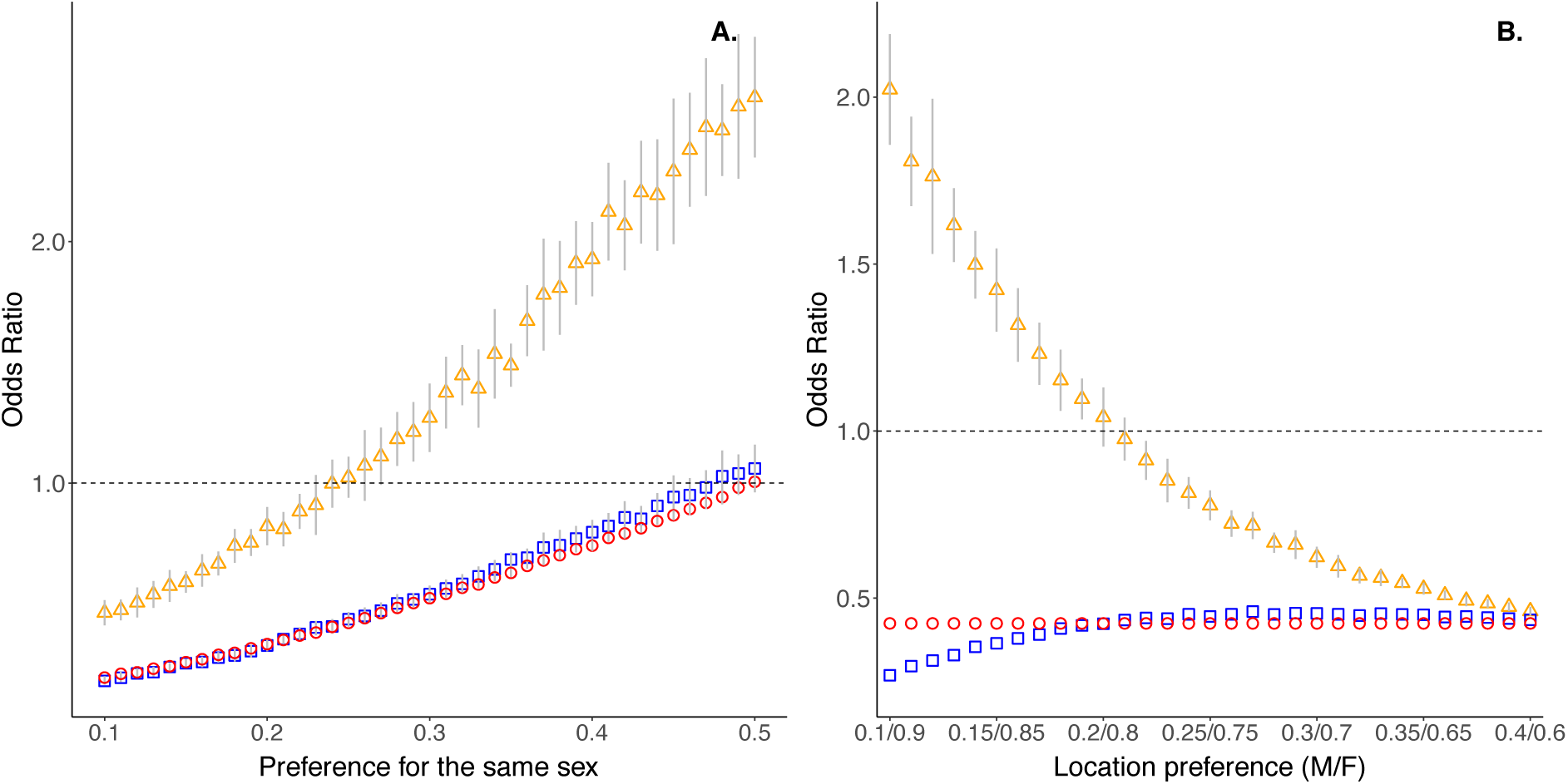
The effect size (odds ratio of associating with the same sex) from the statistical models. Red circles show the effect size for the reference case when males and females share equal location preferences (L_M_= L_F_=0.5). Orange triangles show the effect size of the uncorrected statistical model, and blue squares show the effect size of the model statistically controlling for location overlap. Error bars show standard deviation. Panel A shows results when S_Same_ is varied while S_Different_ = 0.5, L_F_ = 0.8 and L_M_ = 0.2. Panel B shows results when L_F_ and L_M_ are covaried while S_Different_ = 0.5 and S_Same_ = 0.25. These figures show that failing to account for the bias in the statistical model gives incorrect effect sizes, and that this problem can be addressed by controlling for bias in the statistical model.

Unlike the uncorrected model which would rely on permutations to correct confounds only in the p value, the effect of the corrected model can be biologically interpreted. For example, when S_M_=0.7 and S_F_=0.3 and G_M_=0.5 and G_F_=1.0 the corrected mean strength ratio can be interpreted as females being 87% more gregarious than males (reference 82%). This cannot be done for the uncorrected effect size which would be interpreted as males being 4% more gregarious than females. To demonstrate the use of the permutation method with the statistical model, for this example the p values for all 20 runs were significant with p < 0.05 in all cases.

### Simulation 2: Sex assortment and location preferences

The simulation created the desired effect that individuals of a different sex are more likely to associate, while individuals of the same sex are more likely to be in the same location. Over 20 independent simulation runs when males and females had location preferences from the same distribution, we found that two members of a dyad being of the same sex was associated with a decrease in the logit-transformed dyadic association strength of 0.43 (95% CI = 0.42, 0.44). This is the mean (over 20 runs) of the mean posterior expected odds ratio for each simulation, meaning that individuals of the same sex were only 67% less likely to associate than individuals of a different sex at a given sampling event.

For each simulation parameterization our reference/true effect size is from 20 runs of the network under the given parameter values but with location preferences equal for males and females (L_M_= L_F_=0.5). Figure 2 shows the results of our analysis over systematically varied parameter S_Same_, while S_*Different*_ *= 0.5, L*_F_ = *0*.*8* and *L*_M_ = 0.*2* (figure 2a), and over systematically covaried parameters *L*_F_ and *L*_M_ while S_*Different*_ *= 0.5 and* S_*Same*_ = 0.25 (figure 2b). The results show that not accounting for confounding effects in the statistical model produces effect sizes that are very different from the actual effect sizes. Indeed for a wide range of conditions, it actually shows the opposite result to the reference model, in that individuals of the same sex are more likely to associate than individuals of the different sex (points above the dotted line for both panels). However, when we account for sampling in the statistical model, the results are extremely similar to the actual effect sizes.

Unlike the uncorrected model, which would rely on permutations to correct confounds only in the p value, the effect of the corrected model can be interpreted biologically. For example, when S_Same_ *= 0.25.* S_*Different*_ *= 0.5, L*_F_ = *0*.*8* and *L*_M_ = 0.2 odds ratio of associating with the same sex is 0.44, which is extremely close to the true value of 0.43. This cannot be done for the uncorrected effect size which gives a mean odds ratio of 1.02. Under the conditions for this example the p values for all 20 runs were significant with p < 0.05 in all cases.

### Reef Manta Ray Data: Sex differences in gregariousness

In the basic model (strength ∼ sex) the effect size for sex (in this case being male) was −0.45 (SE 0.33), suggesting that males are less gregarious than females. However, in the second model that controlled for observations (strength ∼ sex + number-of-observations) the effect size was −0.07 (SE 0.26) and not statistically significant (GLMQAP permutation test; p = 0.78). Thus, the second model shows that the original effect size was biased by the over sampling of females. The full table of results for both models is provided in the supplementary materials. This offers further support for our finding from the first simulation using data with a more complex structure.

## Discussion

Our study demonstrates that when answering questions related to social preferences, controlling for confounding factors in the fitted statistical model produces biologically meaningful effect size estimates that greatly help the interpretability of the results. Where confounding factors are only controlled for in the null model, as has been typical in animal social network analysis, the only information about the effect comes from the p value and the tail in which significance is found. The latter approach focuses on statistical significance and does not facilitate a careful and balanced assessment of biological importance. As such, it is not desirable to control for confounding factors in permutations alone, and we advocate a return to careful consideration of the statistical model specification.

Our study demonstrates examples of how animal social network analysis can be performed, in the light of recent work highlighting that that datastream permutations do not test the hypothesis that is typically desired with standard regression modelling (Weiss *et al.* 2020). The approach that we have demonstrated a) accounts for confounding factors in the statistical model, b) accounts for the relevant non-independence in the statistical model, and c) uses post-network permutations to further account for non-independence. Thus, we advocate careful thinking about constructing the statistical model to correctly fit the data, including confounding effects. Where possible non-independence structure should also be dealt with in the statistical model, although this can also be dealt with using post-network permutations. Our approach can help facilitate the use of Bayesian analysis, which has possibly been neglected due to the past focus on permutation tests and p values.

If confounding factors are controlled for in the statistical model, then it is unnecessarily to control for them again in the permutations. In some cases no permutations are needed, such as when using dyadic regression with adequate variance/covariance structure (see e.g. our example simulation 2). When the structure to be controlled for is more complex, such as when performing a nodal analysis on a centrality measure, then appropriate permutations can be used, as demonstrated here, such as GLMQAP in the aninet R package for a dyadic analysis (Weiss 2020).

We found that controlling for confounding factors in the statistical dyadic model can help to either mitigate or eliminate the problem, depending on how well these confounding effects can be captured in the model. It often compensates for the extent of the bias, except where there are extreme differences in observations in the first simulation, where it still performs much better than not controlling for confounding factors. Our approach works the same with non-linear approaches such as polynomial regression and GAMs, and we recommend careful examination of predicted model fits and relationships between variables to assess whether a non-linear relationship might be best for the confounding factors.

When analysing with social preferences, there are many situations where confounding factors should be controlled for in the statistical models. Examples of confounding factors include group sizes, location, season, and any known observation bias. When dealing with social preferences, anything that would be controlled for in the null model should ideally be controlled for in the statistical model. Even where the question relates to contacts there will still be occasions where it is appropriate to control for bias in the effect size. For example, here we demonstrated an example of where there is bias for observing certain types of individuals over others, which distorts our observations of the actual interactions (see simulation 1). In situations such as this where differences in observation (or analogously bio-logger performance) may mask the structure of the true social network then controlling for its effect will be important in analysing all animal social networks. However, once confounding factors are controlled for in the statistical model, in most cases it will not be necessary to control for them again in the null model.

Gregariousness is often a key confound for which researchers control (Godde *et al.* 2013). However, in some study systems gregariousness may be a mechanism underlying both the number of observations and the connectedness of individuals in the social network. Where this is the case controlling for the number of observations of each individual in a statistical model can be problematic and including it as a covariate in a model may mask differences in gregariousness between individuals. Another scenario where it is difficult to tease apart competing drivers of association is where social preferences drive use of locations (in addition to spatial preferences impacting association patterns). This is a general problem for networks constructed using data on spatial and temporal co-occurrence that still needs to be resolved, but the key to addressing it is understanding any observation bias and the study system. Simple simulations such as those we have presented here are useful tools in examining the impact of known confounding factors on social preferences prior to any analysis of animal social data, and studying whether they can be adequately controlled for.

## Author Contributions

DWF conceived of the idea and DWF and MNW designed the simulations with input from MJS and DPC; MNW and DWF programmed the simulations; MNW developed and coded the permutations; DWF analysed the results with input from MNW, MJS and DPC; DWF led the writing of the manuscript with input from MNW, MJS and DPC. RJYP collected and provided the reef manta ray data. All authors contributed critically to the final draft and gave final approval for publication.

## Data Accessibility Statement

The manta data will be made accessible on Dydrad, if the paper is accepted for publication.

## Acknowledgements

We thank an anonymous reviewer, Damien Farine, and the editors for comments and suggestions on the manuscript. DWF and DPC acknowledge funding from NERC (NE/S010327/1).

## References

Amrhein, V., Greenland, S. & McShane, B. (2019) Retire statistical significance. Nature, 567, 305–307.

Aplin, L.M., Farine, D.R., Morand-Ferron, J., Cole, E.F., Cockburn, A. & Sheldon, B.C. (2013) Individual personalities predict social behaviour in wild networks of great tits (*Parus major*). Ecology Letters, 16, 1365–1372.

Bejder, L., Fletcher, D. & Brager, S. (1998) A method for testing association patterns of social animals. Animal Behaviour, 56, 719–725.

Brent, L.J.N. (2015) Friends of friends: are indirect connections in social networks important to animal behaviour? Animal Behaviour, 103, 211–222.

Brent, L.J.N., Semple, S., Dubuc, C., Heistermann, M. & MacLarnon, A. (2011) Social capital and physiological stress levels in free-ranging adult female rhesus macaques. Physiology & Behavior, 102, 76–83.

Burnham, K.P., Anderson, D.R. & Huyvaert, K.P. (2011) AIC model selection and multimodel inference in behavioral ecology: some background, observations, and comparisons. Behavioral Ecology and Sociobiology, 65, 23–35.

Butts, C.T. (2016) sna: Tools for Social Network Analysis. R package version 2.4.

Cairns, S.J. & Schwager, S.J. (1987) A Comparison of Association Indexes. Animal Behaviour, 35, 1454–1469.

Carter, A.J., Lee, A.E.G. & Marshall, H.H. (2015) Research questions should drive edge definitions in social network studies. Animal Behaviour, 104, E7–E11.

Chow, C.M., Claxton, S.E. & van Dulmen, M.H.M. (2015) Testing Dyadic Mechanisms the Right Way: A Primer Into Moderated Actor-Partner Interdependence Model With Latent Variable Interactions. Emerging Adulthood, 3, 421–433.

Cohen-Cole, E. & Fletcher, J.M. (2008) Is obesity contagious? Social networks vs. environmental factors in the obesity epidemic. Journal of Health Economics, 27, 1382–1387.

Croft, D.P., Krause, J., Darden, S.K., Ramnarine, I.W., Faria, J.J. & James, R. (2009) Behavioural trait assortment in a social network: patterns and implications. Behavioral Ecology and Sociobiology, 63, 1495–1503.

Croft, D.P., Madden, J.R., Franks, D.W. & James, R. (2011) Hypothesis testing in animal social networks. Trends in Ecology & Evolution, 26, 502–507.

Cumming, G. (2014) The New Statistics: Why and How. Psychological Science, 25, 7–29.

Ellis, S., Franks, D.W., Nattrass, S., Cant, M.A., Weiss, M.N., Giles, D., Balcomb, K.C. & Croft, D.P. (2017) Mortality risk and social network position in resident killer whales: sex differences and the importance of resource abundance. Proceedings of the Royal Society of London B: Biological Sciences, 284.

Farine, D.R. (2017) A guide to null models for animal social network analysis. Methods in Ecology and Evolution, 8, 1309–1320.

Farine, D.R., Aplin, L.M., Sheldon, B.C. & Hoppitt, W. (2015a) Interspecific social networks promote information transmission in wild songbirds. Proceedings of the Royal Society of London B: Biological Sciences, 282.

Farine, D.R., Firth, J.A., Aplin, L.M., Crates, R.A., Culina, A., Garroway, C.J., Hinde, C.A., Kidd, L.R., Milligan, N.D., Psorakis, I., Radersma, R., Verhelst, B., Voelkl, B. & Sheldon, B.C. (2015b) The role of social and ecological processes in structuring animal populations: a case study from automated tracking of wild birds. Royal Society Open Science, 2.

Farine, D.R. & Whitehead, H. (2015) Constructing, conducting and interpreting animal social network analysis. Journal of Animal Ecology, 84, 1144–1163.

Fischer, J., Farnworth, M.S., Sennhenn-Reulen, H. & Hammerschmidt, K. (2017) Quantifying social complexity. Animal Behaviour, 130, 57–66.

Fisher, D.N., Ilany, A., Silk, M.J. & Tregenza, T. (2017) Analysing animal social network dynamics: the potential of stochastic actor-oriented models. Journal of Animal Ecology, 86, 202–212.

Garcia, R.L., Kenny, D.A. & Ledermann, T. (2015) Moderation in the actor-partner interdependence model. Personal Relationships, 22, 8–29.

Godde, S., Humbert, L., Cote, S.D., Reale, D. & Whitehead, H. (2013) Correcting for the impact of gregariousness in social network analyses. Animal Behaviour, 85, 553–558.

Halsey, L.G. (2019) The reign of the p-value is over: what alternative analyses could we employ to fill the power vacuum? Biology Letters, 15.

Halsey, L.G., Curran-Everett, D., Vowler, S.L. & Drummond, G.B. (2015) The fickle P value generates irreproducible results. Nature Methods, 12, 179–185.

Hamede, R.K., Bashford, J., McCallum, H. & Jones, M. (2009) Contact networks in a wild Tasmanian devil (*Sarcophilus harrisii*) population: using social network analysis to reveal seasonal variability in social behaviour and its implications for transmission of devil facial tumour disease. Ecology Letters, 12, 1147–1157.

Hoff, P.D., Raftery, A.E. & Handcock, M.S. (2002) Latent Space Approaches to Social Network Analysis. Journal of the American Statistical Association, 97.

Ilany, A., Booms, A.S. & Holekamp, K.E. (2015) Topological effects of network structure on long-term social network dynamics in a wild mammal. Ecology Letters, 18, 687–695.

Kenny, D.A. (1996) Models of non-independence in dyadic research. Journal of Social and Personal Relationships, 13, 279–294.

Kenny, D.A. (2018) Reflections on the actor-partner interdependence model. Personal Relationships, 25, 160–170.

Krackhardt, D. (1987) Qap Partialling as a Test of Spuriousness. Social Networks, 9, 171–186.

Krackhardt, D. (1988) Predicting with Networks - Nonparametric Multiple-Regression Analysis of Dyadic Data. Social Networks, 10, 359–381.

Nakagawa, S. & Cuthill, I.C. (2009) Effect size, confidence interval and statistical significance: a practical guide for biologists. (vol 82, pg 591, 2007). Biological Reviews, 84, 515–515.

Newman, M.E. (2004) Analysis of weighted networks. Phys Rev E Stat Nonlin Soft Matter Phys, 70, 056131.

Perryman, R.J.Y., Venables, S.K., Tapilatu, R.F., Marshall, A.D., Brown, C. & Franks, D.W. (2019) Social preferences and network structure in a population of reef manta rays. Behavioral Ecology and Sociobiology, 73.

R Core Team (2018) R: A language and environment for statistical computing. R Foundation for Statistical Computing, Vienna, Austria.

Ramos-Fernandez, G., King, A.J., Beehner, J.C., Bergman, T.J., Crofoot, M.C., Di Fiore, A., Lehmann, J., Schaffner, C.M., Snyder-Mackler, N., Zuberbuhler, K., Aureli, F. & Boyer, D. (2018) Quantifying uncertainty due to fission-fusion dynamics as a component of social complexity. Proceedings of the Royal Society of London B: Biological Sciences, 285.

Rushmore, J., Caillaud, D., Matamba, L., Stumpf, R.M., Borgatti, S.P. & Altizer, S. (2013) Social network analysis of wild chimpanzees provides insights for predicting infectious disease risk. Journal of Animal Ecology, 82, 976–986.

Silk, J.B., Beehner, J.C., Bergman, T.J., Crockford, C., Engh, A.L., Moscovice, L.R., Wittig, R.M., Seyfarth, R.M. & Cheney, D.L. (2009) The benefits of social capital: close social bonds among female baboons enhance offspring survival. Proceedings of the Royal Society of London B: Biological Sciences, 276, 3099–3104.

Silk, M.J., Croft, D.P., Delahay, R.J., Hodgson, D.J., Boots, M., Weber, N. & McDonald, R.A. (2017) Using social network measures in wildlife disease ecology, epidemiology, and management. Bioscience, 67, 245–257.

Silk, M.J. & Fisher, D.N. (2017) Understanding animal social structure: exponential random graph models in animal behaviour research. Animal Behaviour, 132, 137–146.

Weiss, M.N. (2020) Aninet R Library: Statistical models for animal social networks.

Weiss, M.N., Franks, D.W., Brent, L.J., Ellis, S., Silk, M.J. & Croft, D.P. (2020) Common datastream permutations of animal social network data are not appropriate for hypothesis testing using regression models. bioRxiv, doi: 10.1101/2020.04.29.068056.

Weiss, M.N., Franks, D.W., Croft, D.P. & Whitehead, H. (2019) Measuring the complexity of social associations using mixture models. Behavioral Ecology and Sociobiology, 73.

Whitehead, H. (2008) Analyzing Animal Societies: Quantitative Methods for Vertebrate Social Analysis. University of Chicago Press.

Whitehead, H. & James, R. (2015) Generalized affiliation indices extract affiliations from social network data (vol 6, pg 836, 2015). Methods in Ecology and Evolution, 8, 1645–1645.

