## supplementary materials for "Calculating effect sizes in animal social network analysis"

**Supporting information for: Animal Social Network Analysis Has an Effect Size Problem**

Below are the model outputs for the manta analysis.

A) The uncorrected regression model.

| | Estimate | Std. Error | t value |
|---|---|---|---|
| (Intercept) | 5.6201 | 0.2034 | 27.637 |
| sex(M) | -0.4452 | 0.3321 | -1.341 |

B) The corrected regression model.

| | Estimate | Std. Error | t value | Pr(Permuted) |
|---|---|---|---|---|
| (Intercept) | 5.48081939 | 0.15956309 | 34.3489181 | NA |
| sex(M) | -0.07372855 | 0.26286649 | -0.2804791 | 0.797202797 |
| totalSeen meanCentred | 0.14028677 | 0.01658053 | 8.4609338 | 0.001998002 |
